# Successful establishment of an exogenous gene expression system in *Chlamydomnas* chloroplast based on viral replication elements

**DOI:** 10.1101/2024.11.26.625556

**Authors:** Yanan Jiang, Guiying Zhang, Song Wang, Ali Raza, Xiang Liu, Chunli Guo, Sha Wu, Jixu Luo, Yihao Lie, Yingqi Liu, Bin Jia, Zhangli Hu

## Abstract

Currently, transformation in unicellular green alga (*Chlamydomonas reinhardtii*) chloroplast is hindered by a low yield of target products, possibly due to the heterogeneity of the chloroplast genome. In this study, an expression platform was established in *C. reinhardtii* by leveraging replicative elements from beet curly top geminivirus, which led to a high level of heterologous gene expression, as previously reported in *Nicotiana tabacum*. Platform performance was verified by the successful expression of a linearized *mvenus* cassette, achieving a higher expression level than that obtained using the homologous-recombination-based approach. This study provides a new strategy in microalgae synthetic biology and opens new avenue for accumulating valuable bioproducts at high concentrations.

## 1 Introduction

Microalgae have been considered promising organisms as they, in contrast to bacteria, can synthesize bioproducts via photosynthesis (Crozet et al., 2018). Microalgae-derived biochemicals include lipids, protein pigments, carbohydrates, vitamins, antioxidants, and unsaturated fatty acids (Chew et al., 2017). *Chlamydomonas reinhardtii*, as the best understood green algae, draws significant attention due to its superiority in biosynthesizing bioactive substances containing recombinant protein and small molecules (Rasala et al., 2015). Currently, *C. reinhardtii* has been used as cell factory to produce carbohydrates, fatty acids, functional proteins, pigments, hormones, vaccines, and antibodies (Masi et al., 2023). *C. reinhardtii* also serves as an excellent model for studying photosynthesis and flagella-mediated movement. Its consumption is considered safe, as it has been classified as GRAS. *C. reinhardtii* is devoid of endotoxins and viral or prion contaminants, which is particularly beneficial for the production of therapeutic proteins (Stoffels et al., 2019). Moreover, the three sets of genomes (nuclear, chloroplastic, and mitochondrial genomes) of *C. reinhardtii* have been sequenced, and transformations have been established for all three sets of genomes even though transcription and translation vary (Merchant et al., 2007). Synthesis of recombinant proteins with *C. reinhardtii* offers multiple advantages compared to other chassis, such as mammalian cells, bacteria, and yeast. For example, large-scale cultivation of *C. reinhardtii* can be carried out at a low cost, and it is not susceptible to contamination; *C. reinhardtii* can process certain proteins that bacteria and fungi cannot fold properly (Specht et al., 2010; Chebolu et al., 2009; Franklin et al., 2004). Therefore, *C. reinhardtii* has been developed as an excellent platform for functional proteins, metabolism-related enzymes, high-value products, and proteins for pharmaceutical purposes. Class-II metallothionein, GFP, HBcAgII, luciferase, and ZEP have been successfully expressed in *C. reinhardtii* (Zhang et al., 2021).

*C. reinhardtii* chloroplast is an excellent platform for synthetic biology, as it is a relatively small organelle, and contains most of the photosynthesis-related genes. Chloroplast enables *C. reinhardtii* to convert natural light energy into chemical energy (O’Neill et al., 2012). Compared with nuclear production of recombinant proteins in *C. reinhardtii*, chloroplast-based production has confirmed to be more promising and advantageous in several aspects. Firstly, there is no gene silencing in chloroplast, which can increase gene expression (Bock et al., 2007). Moreover, chloroplast is a prokaryotic expression system, allowing multiple genes to be regulated by a single promoter (Bock et al., 2007; Rymarquis et al., 2006). Finally, recombinant proteins expressed in chloroplasts are not glycosylated, which is particularly important for the production of monoclonal antibodies, as the majority of therapeutic human monoclonal antibodies require no glycosylation (Franklin et al., 2005; Dove et al., 2002). The recombinant proteins produced in *C. reinhardtii* chloroplasts, such as green fluorescent protein (GFP), human metalthianine-2, and aminoglycoside adenine transferases (Mayfield et al., 2007).

At present, transformation of foreign DNA into the *C. reinhardtii* chloroplasts relies on homologous recombination, which can accurately insert foreign fragments into the chloroplastic genome (Esland et al., 2018). The established approaches include particle bombardment, glass bead and electroporation transformation (Shamriz et al., 2016). However, particle bombardment suffers from high cost, while the glass bead method is only suitable for *C. reinhardtii* species without a cell wall. The electroporation method is not commonly used for chloroplastic transformation (Ma et al., 2022). The existence of more than 80 copies of the chloroplast genome makes chloroplastic transformation even more challenging (Rosales-Mendoza et al., 2012). Therefore, it takes a long time to obtain a genetically stable transformant with genome containing all endogenous sequnce replaced by foreign sequences. Additionally, screening pressure should always be maintained; otherwise, foreign genes may be replaced by wild-type gene fragments in subsequent subcultures, resulting in the loss of foreign genes. The construction of gene expression vectors for chloroplast transformation of *C. reinhardtii* is complicated and expensive, and the regulatory mechanism of gene expression remain unclear, leading to a low expression of exogenous proteins. Therefore, a novel and efficient method is urgently needed to increase the expression level. As introduced previously, in contrast to insertion into the chloroplastic genome via homologous recombination, replication elements from germini virus can achieve independent circular DNA molecules and potentially accumulate DNA sequences of interest to a high level (Lozano-Durán et al., 2016). Rep protein plays a vital role in the plant viral duplication strategy (Lozano-Durán et al., 2016). It creates a break in the intergenic regions and initiates a rolling circle replication and circularization of the new DNA molecules to achieve a high copy number in host cells (Stenger et al., 1991). This virus-derived strategy was proven successful in editing the plants genome but its exploitation in algal chloroplasts has not been evaluated yet (Jakubiec et al., 2021).

In this work, we constructed a novel chloroplast transformation strategy in *C. reinhardtii* for the first time, leveraging a germinivirus replication protein (Rep) and its replication element (vor) (Jakubiec et al., 2021). A chassis alga was constructed by expressing Rep in *C. reinhardtii*, and mVenus was used to verify the constructed system. Our results show that episomal DNA containing *mvenus* cassette was formed, and mVenus was successfully expressed in *C. reinhardtii* chloroplasts with Rep and vor. In short, our work provides a new and efficient method for synthesizing bioproducts in chloroplosts of *C. reinhardtii*.

## 2 Material and methods

### 2.1 Algae and cultivation

Wild-type *C. reinhardtii* (CC-125) was purchased from the Freshwater Algae Culture Collection at the Institute of Hydrobiology (FACHB), National Aquatic Biological Resource Center, China, and cultured at 25 □ under 90 μmol/m^2^/s light intensity using TAP medium.

### 2.2 Plasmids preparation

#### 2.2.1 Construction of expression vectors

The sequence of Rep in Rep-aphvIII vector from tobacco beet curly top geminivirus (BCTV) (Jakubiec et al., 2021). The *rep* required for this experiment was chemically synthesized by GenScript (Nanjing, China) with nuclear codon optimization according to the codon bias of *C. reinhardtii* chloroplast genome. The plasmid (Rep-aphvIII) contains the CDS of the *rep* (Jakubiec et al., 2021), an HSP70Ap/RBCS2p fusion promoter/terminator, a chloroplast transit peptide psaD (Fischer et al., 2001), a Strep-Tag II protein tag, an *aphvIII* resistance gene cassette, as well as *E. coli amp* resistance cassette. The genes (*mvenus* and *vor)* required for the experiment were synthesized after optimization. The mVenus sequence (AAZ65844.1) and the regulatory element *vor* sequence (NC_001412.1) from BCTV were retrieved from NCBI. The plasmid containing mVenus protein (vor-mvenus-vor vector) includes the CDS of *mvenus*, regulated by psaD promoter and psaD terminator, an *aada* resistance expression cassette, and *E. coli amp* resistance cassette. The *mvenus* cassette and the *aada* resistance cassette were flanked by two reverse repeating *vor* sequences. The plasmid of mVenus protein (p322-mvenus expression vector) contains CDS of *mvenus*, regulated by psaD promoter and psaD terminator, *aada* resistance expression cassette as well as *E. coli amp* resistance cassette. The *mvenus* cassette and *aada* resistance cassette were flanked by homologous sequences of *psaa* gene in the chloroplast genome of *C. reinhardtii*.

### 2.3 Transformation

#### 2.3.1 Nuclear transformation

A single clone of *C. reinhardtii* (CC-125) from a plate grown in TAP medium at 25 °C under standard light (90 μmol/m^2^/s) for seven days, was inoculated into 50 mL TAP liquid medium at 25 □ under constant light condition (90 μmol/m^2^/s, 24 hours light), and cultured in a shaking incubator for four days. Then, one mL of the *C. reinhardtii* culture medium was inoculated into a fresh 100 mL liquid TAP medium and cultured in a shaker for three to four days. The *C. reinhardtii* cells in exponential phase were collected by centrifugation at 1200 *g* for five minutes and resuspended to a cell density of 10^8^ cells/mL. Electroporation was performed with GenePulser^®^II with a voltage of 500 V, capacitance 50 μF, and resistance of 800 Ω. The electroporation cuvette had a gap distance of four mm. After incubation at low light for 24 h, the transformed algae were spread on TAP solid plates containing five μg/mL parommomycin at 22-25 □, under continuous light (90 μmol/m^2^/s), and thecolionies were appeared after 10-14 days.

#### 2.3.2 Chloroplastic transformation by electroporation

The culture conditions of *C. reinhardtii* are as follows: a single colony was picked from the *C. reinhardtii* plate and inoculated into 100 mL TAP liquid medium at 25 □ under light conditions (about 90 μmol/m^2^/s). After six days of cultivation, one mL of *C. reinhardtii* culture was inoculated into 200 mL fresh liquid TAP medium and cultured in a shaker, and the electroporation transformation experiment was carried out within 24 h of culture. During this period, 200 mL of fresh algal liquid was pre-cooled on ice, and filtered Tween 20 was added to a final concentration of 1/2000 (v/v). The suspension was centrifuged at 2000 g at four □ for five minutes. The collected *C. reinhardtii* cells were washed with cold ddH_2_O and then re-suspended with two mL ddH_2_O, reaching a final density of 10^8^ cells/mL. Algal cells (100 μ□), 100 μL 50% PEG3350, 20 μL salmon extract DNA (10 mg/mL) and one μg plasmid DNA were mixed evenly. Electroporation was performed using the following parameters: capacitance (25 μF), voltage (220 V), resistance (∞ Ω) and cuvette (one mm). After the electroporation pulse was completed, the suspension was recovered in 10 mL TAP for 24 hours at 25 °C in the dark. After recovery, the transformed solution was centrifuged, resuspended to one mL, and 200 μL aliquots were plated on TAP solid medium containing five μg/mL paromomycin and 50 μg/mL spectinomycin.Transformants were cultured under continuous light (90 μmol/m^2^/s) at 22-24 □ for six-eight days.

#### 2.2.3 Chloroplastic transformation with biolistic bombardment

The *mvenus* expression cassette was integrated into the *C. reinhardtii* chloroplast genome by homologous recombination using the vector p322-mvenus with a homologous sequence of *psaa* gene. The p322-mvenus plasmid was extracted and concentrated to one μg/μL. Suspension of CC-125 (50 mL) was centrifuged at room temperature at 700 *g* for five minutes, with cells suspended to a density of 10^8^ cells/mL, and the concentrated algal suspension was evenly coated in the middle of TAP solid plates. Five μL DNA (one μg/μL), 50 μL of 2.5 M CaCl_2_, and 20 μL of 0.1 M spermidine to 50 μL (three mg) of microcarrier solution successively were added while vortexing. The mixture was centrifuged at 700 *g* for five minutes; the supernatant was discarded in the super-clean workbench, washed with 500 μL 70% ethanol and then 500 μL of ethanol successively, and finally resuspended in 100 μL ethanol. During the gene gun transformation process, the embedded microvarriers are placed on the carrier membrane. Susequently, 10 μL of the embedded microcarriers onto the center of the carrier membrane and allow it to sit for 10 mintues to enable ethanol evaporation. The bombardment was carried out by the Biolistic PDS-1000/He particle delivery system (BIO-RAD, California, USA) with 1350 psi rupture disks at a distance of nine centimeters. After the bombardment, the plates were incubated in darkness for 24 h to allow for revival and then observed for one to two weeks before selecting positive transformants.

### 2.3 Identification of transformants

The colonies of Rep-aphvIII, vor-mvenus-vor, and p322-mvenus of *C. reinhardtii* on selective plates were picked and transferred to new solid TAP plates containing paromomycin and placed under continuous light (90 μmol/m^2^/s) at 25 □. Colonies were further inoculated into one mL TAP liquid culture medium in 1.5 mL microcentrifuge tubes. After three to four days cultivation, DNA was extracted using a universal rapid extraction DNA kit (081001, Beibei Biotechnology, China), following the manual guidelines. Lastly, PCR was performed to verify the transformation.

### 2.4 Western blot

To verify the successful expression of mVenus in the transformed strain, western blot quantitative analysis was performed. Protein quantification was conducted using the BCA Protein Assay (GK10009, GLPBIO, USA), following the kit instructions. eGFP Monoclonal Antibody (MA1-952, Thermo Fisher Scientific, USA) was used as the primary anti-GFP Antibody, and beta Tubulin Monoclonal Antibody (AA10, Thermo Fisher Scientific, USA) served as the internal reference protein. The positive control was the algal strain extract containing the relevant target genes, while the negative control was the total protein extract of wild-type (CC-125). The specific steps are as follows: 200 mL algal suspension was subjected to freeze-thaw cycles to lyse the cells and extract total protein. Protein was precipitated using acetone pre-cooled at -20 °C. After removing the supernatant, PBS was added to dissolve the protein precipitate. SDS-PAGE was performed using SDS-PAGE protein sample buffer (HR0368, Biolab, China), gel (F15412MGel, ACE, China), and TRIS-Glycinate electrophoresis solution (BR0002-01, ACE, China). Proteins were then transferred onto a 0.22 μm PVDF membrane (IPVH00010, MerckMillipore, Germany). The membrane was sealed at room temperature for one h with QuickBlock™ Western sealing solution (P0252, Beyotime Biotechnology, China) before incubation with primaryand secondary antibodies (Beyotime Biotechnology, China). Chemiluminescence was detected using BeyoECL Star (P0018AS, Beyotime Biotechnology, China) with the Tanon 5200 imaging system (Tanon, China).

### 2.5 Southern blot

The obtained Rep-mvenus transformed strain was selected from the plate, and the monoculture of *C. reinhardtii* was grown in fresh 100 mL TAP for four days as inoculum. Starter culture (five mL) was added to 200 mL TAP under continuous light (90 μmol/m^2^/s) at 25 □ until the cells reachers the logarithmic growth stage. Genomic DNA was extracted using Plant DNA Kit (D3488-02, Omega Bio-Tek, USA). The extracted DNA was digested with 10 µg *EcoRI* restriction enzymes in a total volume of 50 µL at 37 L for six h. DNA fragments were separated by 0.8% agarose gel electrophoresis and then transferred to a nylon membrane. Southern blot analysis was performed using digoxigenin DNA Labeling and Detection Kit (11093657910, Roche, Switzerland), following the manufacturer’s instructions. Primer F-114/R-637 was used to design a specific probe for the sequence near *mvenus* was designed using the primers F-114 (5*’*-CACCTACGGCAAGCTGACCCT-3*’*) and R-637 (5*’*-CCATGTGATCGCGCTTCTCGTT-3*’*).

### 2.6 Confocal laser scanning microscopy observation

An ultra-high resolution confocal laser scanning microscope (LSM 710 NLO, ZEISS, Germany) was used for observation. The fluorescence microscope parameters were set as follows: the excitation wavelength for mVenus was 488 nm, and the emission wavelength was 509 nm. The excitation and emission wavelengths for chlorophyll a were 433 and 664 nm, respectively.

### 2.7 Statistical data analysis

Biological triplicates were prerformed in every measurement and Students’ *t* test was applied to evaluate significance.

## 3 Results and discussion

### 3.1 Establishment of a chassis strain for chloroplastic episome expression

Previous studies have shown that a novel chloroplast expression platform can be developed in tobacco using virus-related elements (Jakubiec et al., 2021). In this study, a Rep-aphvIII transformation vector harboring *rep* gene of BCTV and the paromomycin (*aphvIII*) resistance gene cassette was constructed (Fig. 1B) to serve as chassis cells. A chloroplast targeting sequence (PsaD-target) was fused with the *rep* gene, which is expressed in the nucleus, to guide it into the chloroplast. The vor-mvenus-vor plasmid, transformed via electroporation, contained a *mvenus* expression cassette driven by PsaD promoter, flanked by complementary *vor* sequences. Rep was able to recognize one of the *vor* sequences on the vor-mvenus-vor vector and initiate replication by generating a niche. Circularization was achieved via reverse complementation between the two *vor* sequences (Fig. 1A). Western blot analysis confirmed the successful expression of Rep in *C. reinhardtii*. Of the five positive strains, the strain with the highest Rep expression (Rep-4) was selected as chassis for epsiome expression in *C. reinhardtii* chloroplast (Fig. 1C).

**Figure 1.**
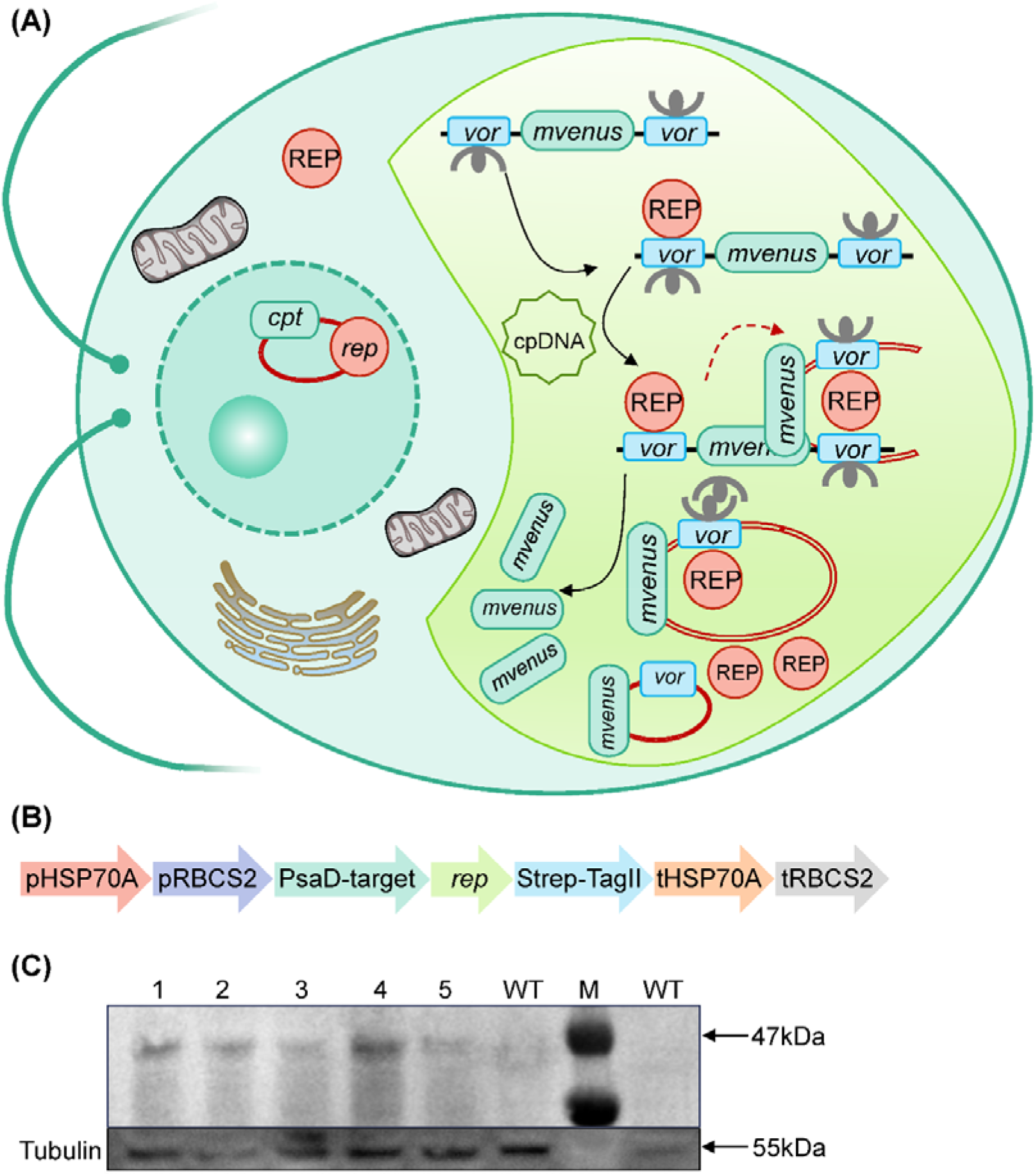
Establishment of the chassis cell and underpinned mechanism. **(A)** cpt: chloroplast targeting peptide sequence; *vor* sequence: sequence specifically recognized by the REP protein from beet curly top bipyramidal virus (BCTV); *mvenus*: gene of yellow fluorescent protein; *rep*: gene of BCTV. **(B)** pHSP70A/pRBCS2 is the promoter; PsaD-target: chloroplast trageting peptides; rep is the *rep* gene of BCTV; Strep-TagII is the protein tag; tHSP70A/tRBCS2 is the terminator. **(C)** Lane 1-5: transformants of Rep; WT: wild type; Strep-Tag II is protein tag (47 kDa); Tubulin (55 kDa) serves as reference.

*C. reinhardtii* has emerged as a promising cell factory for the production of diverse bioactive substances. The chloroplast of *C. reinhardtii* has been acknowledge as a platform where transgene can be incorporated into chloroplast genome via homologous recombination (Dyo et al., 2018) (Doron et al., 2016). Nevertheless, due to the muti-copy characteristics of chloroplast genome, reaching acceptable and stable expression of exogenous proteins requires extensive homogenization, which remains both challenging and time-consuming.. In 2021, a minichromosome strategy in N. tabacum by leveraging elements from BCTV was developed to lead high-level transgene expression and effectively circumventing the issue of heterogenization (Jakubiec et al., 2021). In this study, we further verified these virus-derived elements in a unicellular photosynthetic chassis for synthetic biology. The advantages of the episomal platform over traditional homologous recombination are clearly demonstrated by the higher accumulation of mVenus in *C. reinhardtii* chloroplast. This approach can be extended to synthesising other exogenous proteins in future applications.

### 3.2 Verification of the episome expression system

To verify whether the episome system functioned properly in *C. reinhardtii*, we constructed the vor-mvenus vector containing the mVenus yellow fluorescent protein, regulated by PsaD promoter and PsaD terminator, flanked by reverse repeats of *vor* sequences on both sides of the *mvenus* expression cassette. These *vor* sequences can be specifically recognized by Rep proteins (Fig. 2A). This vector was transformed into the chloroplasts of the chassis (Rep-4), obtained in the previous step, via electroporation. The Rep protein, guided into plastic, could specifically recognize the *vor* sequence in the vector and initiate replication (Fig. 1A).

**Figure 2.**
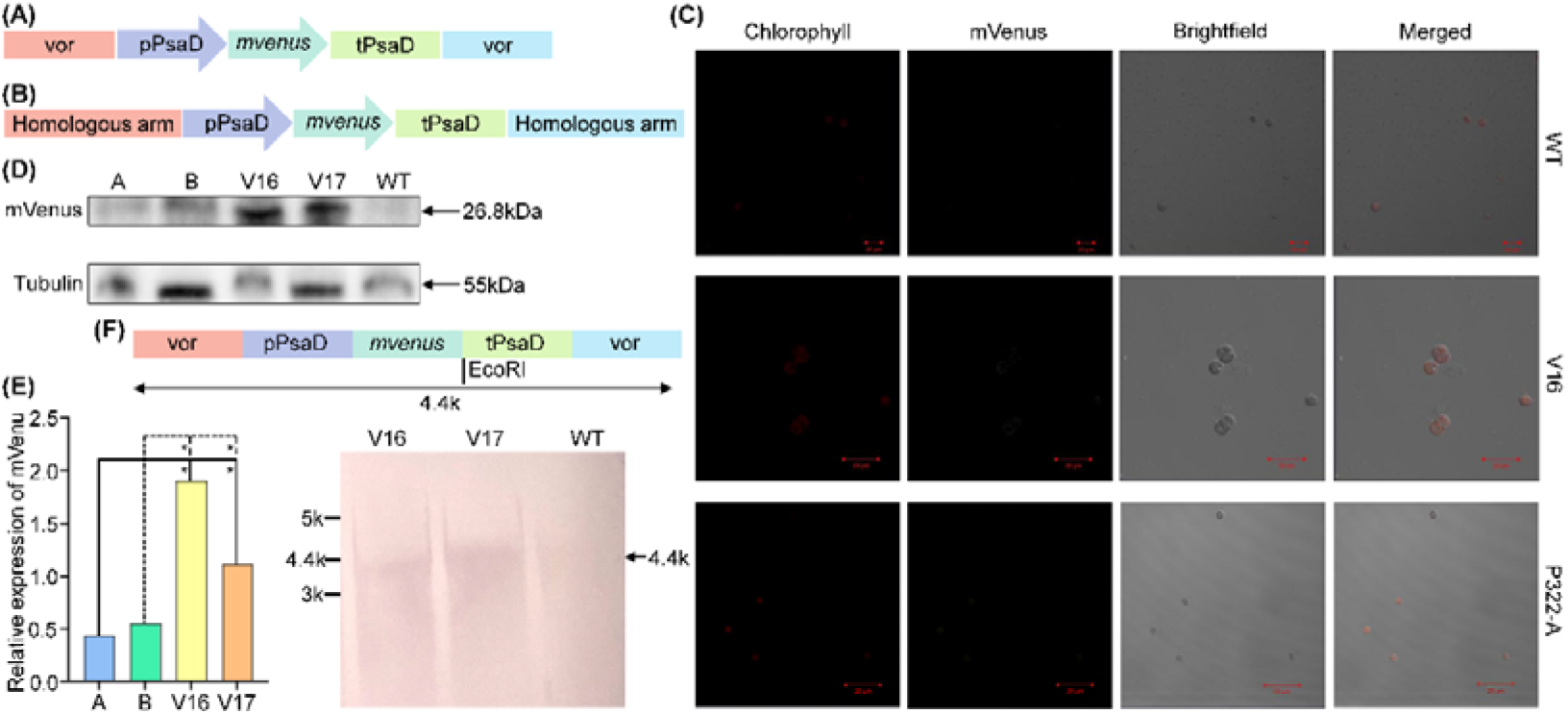
Verification of the effectiveness of the chloroplastic episome system. **(A)** *vor* sequence: sequence specifically recognized by the REP from BCTV; *mvenus*: gene of yellow fluorescent protein; *rep*: gene of BCTV; pPsaD and tPsaD are promoter and terminator, respectively. **(B)** Transformation vector into chloroplast via homologous recombination. **(C)** Confocal microscopic images of transformants with laser scanning confocal microscope. WT: wild type; V16: the transformants obtained with the episome system; P322-A: transformant obtained by homologous recombination. **(D)** Western blot. Hybridization was performed using GFP antibody. and a 26.8 kDa band was observed in the transformants, WT is the wild-type *C. reinhardtii* (CC-125), V16/V17 are the transformants obtained by the episome system, and A/B are the transformants obtained by homologous recombination. **(E)** Gray-scale analysis of western blotting results. (F) Southern blot

Moreover, we constructed the p322-mvenus vector, in which the *mvenus* expression cassette was flanked by homologous arms of the *C. reinhardtii* chloroplast genome. (Fig. 2B). Then, we compared the fluorescence of mVenus in both systems, as well as its expression at the transcriptional level. Confocal fluorescence microscopy was performed on the obtained transformants, suggesting that the yellow fluorescent protein could be observed when excited at 488 nm with an emission wavelength of 509 nm (Fig. 2C). The results displayed that the fluorescence intensity of mVenus expressed through the episome system (vor-mvenus-V16) was higher than that achieved by homologous recombination (p322-mvenus-A).

Western blot analysis using GFP antibodies was performed on two transformants from each approach. With equal protein loading, a band at 26.8 kDa was observed in the transformants obtained through the episome system. In comparison, the A/B transformants from the homologous recombination approach displayed a relatively weaker band at 26.8 kDa (Fig. 2D). Image J was used to perform gray -scale analysis of bands in Western blot. The results showed that the protein expression level of mVenus in transformants V16 and V17 obtained by episomal system was significantly higher than that of transformants A/B obtained by homologous recombination (p < 0.05) (Fig. 2E). This analysis suggest that the amount of fluorescent proteins expressed by homologous recombination was lower than that expressed in the episome form, consistent with the results of confocal fluorescence microscopys. This analysis suggest that the amount of fluorescent proteins expressed by homologous recombination was lower than that expressed in the episome form, consistent with the results of confocal fluorescence microscopys.

Additionally, we performed Southern blot analysis on the transformants obtained through the episome system to verify that mVenus expression is indeed occurring in a circularized plasmid via Rep protein recognition. A probe was designed at the *mvenus* gene, and if the expression of *mvenus* occurred in a loop through the episome, the probe would hybridize with a single 4.4 kb band after digestion by *EcoRI* enzyme (Fig. 2F). The results showed that only a 4.4 kb band was observed in the transformants, confirmating that the episome system proposed in this experiment functioned appropriately in *C. reinhardtii*.

## 4 Conclusion

In short, this study established a chloroplastic expression platform in the model *C. reinhardtii* by leveraging virus-derived replication elements. The platform’s performance was verified by the higher accumulation of mVenus as reporter and heterologous expression of carotenoid. Therefore, this novel episomal platform delivers a more efficient and effective alternative to traditional chloroplastic expression systems for producing heterologous proteins and bioproducts via homologous recombination.

